# ILC2 cells promote lung cancer and accumulate in tumors concomitantly with immune-suppressive cells in humans and mice

**DOI:** 10.1101/2023.05.04.539356

**Authors:** Ilham Bahhar, Zeynep Eş, Oğuzhan Köse, Akif Turna, Mehmet Zeki Gunluoglu, Aslı Çakır, Deniz Duralı, Fay C. Magnusson

## Abstract

It is now clear that group 2 innate lymphoid cells (ILC2) play crucial and sometimes opposing roles in the lung, such as restoring barrier function and integrity after viral infections or, on the contrary, exacerbating inflammation and tissue damage in allergic asthma. However, their role in lung cancer is still unclear. Here, we report that human non-small cell lung cancer patients bear increased frequencies of ILC2s in tumors, normal lung tissue and peripheral blood (PB) as compared to PB from healthy donors (HDs). Frequencies of Foxp3^+^ regulatory T cells were also increased in NSCLC patients, concomitantly with ILC2s. In mice bearing heterotopic lung cancer, adoptive transfer of ILC2s led to increased tumor growth and reduced survival. The frequencies of monocytic myeloid-derived suppressor cells (M-MDSCs) were found to be increased in the tumors of mice that received ILC2s as compared to controls. Overall, our results indicate that ILC2 cells play a pro-tumoral role in lung cancer potentially by recruiting immune-suppressive cells to the tumors.

## INTRODUCTION

Group 2 innate lymphoid cells (ILC2s) are members of the newly discovered family of innate immune cells that play key roles within barrier tissues. They rapidly respond to tissue insult or injury by releasing interleukin-4 (IL-4), IL-5 and IL-13 and they promote a type 2 adaptive immune response (Vivier et al. 2018). ILC2 cells are known to be enriched in both human and mouse lung, and to be activated by IL-25 and IL-33 (Cheng et al. 2017).

Lung cancer is the leading cause of cancer-related deaths worldwide, with non-small cell lung cancer (NSCLC) being the most common (80%) lung cancer subtype (Siegel et al. 2022).

The role of ILC2 cells in cancer is highly controversial with some studies showing they are protumorigenic, while others report the opposite (Magnusson and Bahhar 2023; Trabanelli et al. 2019). In recent years, the number of publications reporting on the role of ILC2 cells in cancer contexts have grown exponentially (Trabanelli et al. 2019). Nevertheless, a clear role for ILC2 cells in cancer development and metastasis is still not established, underlying how tissue-specific microenvironmental cues may lead to different outcomes. Moreover, recent evidence has called for a distinction between IL-33-activated ILC2 cells and IL-25-activated ILC2 cells as two different subsets of ILC2s with potentially distinct functions (Y. Huang and Paul 2016).

Studies of ILC2 cells in the context of lung cancer are sparse, both in mice and humans. In humans, while ILC2 cells were found to be enriched in NSCLC patients as compared to healthy donors (HDs), suggesting a pro-tumoral role for ILC2 cells (Wu et al. 2017; Shen et al. 2021), seemingly conflicting results were also reported, as ILC2 frequencies were found to be reduced in tumors as compared to normal lung tissue of NSCLC patients (Carrega et al. 2015). The conflicts may be due to differing populations taken as baseline reference, therefore additional and more comprehensive studies comparing the frequencies of ILC2 cells in various tissues of NSCLC patients and HDs are needed.

In mice, IL-33-activated ILC2 cells enhanced anti-tumor immunity in a model for primary and metastatic lung tumor (Saranchova et al. 2018). Conversely, they facilitated metastasis to the lung by suppressing natural killer (NK) cells (Schuijs et al. 2020). Such opposing functions for IL-33-activated ILC2 cells have also been reported in other cancer types such as models for pancreatic cancer (Moral et al. 2020). In a mouse model for spontaneous colorectal cancer (CRC), the IL-25-ILC2 axis was found to be pro-tumoral (Jou et al. 2022), while inhibiting IL-25 signaling or ILC2 cells in an induced model of CRC led to an increased number of tumors (Q. Huang et al. 2021; Thelen, Green, and Ziegler 2016). These discrepancies highlight the importance of tissue-specific tumor microenvironments as well as their associated ILC2-activating signals when studying the role of ILC2 cells in cancer. To date, the role of IL-25-activated ILC2 cells in a murine model of lung cancer remains unexplored.

Here, we studied the role of ILC2 cells in lung cancer both in humans and in mice. We show that ILC2 cells are enriched in tumor tissue, adjacent normal lung tissue and PB of NSCLC patients as compared to HDs. Furthermore, we found that ILC2 cells accumulated in tumors as compared to adjacent normal lung tissue. Concomitant with the increase in ILC2 cells, we also found an increase in immunosuppressive Foxp3^+^ regulatory T cells (Tregs) in tumor tissue, adjacent normal lung tissue and mediastinal lymph nodes (LNs) of NSCLC patients as compared to HDs. Adoptive transfer of IL-25-activated ILC2 cells in lung cancer-bearing mice led to increased tumor burden and reduced survival. Frequencies of monocytic myeloid-derived suppressor cells (M-MDSCs) were increased in tumors of mice that received ILC2 cells as compared to controls, which may be due to the high production of IL-13 and IL-4 by ILC2 cells. Taken together, our findings highlight a pro-tumoral function for ILC2 cells in NSCLC both in humans and in mice and suggest that these cells may constitute a promising therapeutic target in NSCLC.

## MATERIAL AND METHODS

### Patients and Healthy donors

Peripheral blood, tumor sample, along with an adjacent non-invaded tissue sample and mediastinal lymph node sample were collected from 40 patients with NSCLC undergoing surgical resection at Cerrahpasa Hospital and Medipol Hospital in Istanbul. Prior to sampling, none of the patients had received chemotherapy or radiation therapy. All subjects gave written informed consent to participate in this study. The study protocol was approved by Istanbul Medipol University Ethics Committee. Details of the clinicopathological characteristics of these patients are summarized in Supplementary Table S1. As a control, peripheral blood samples were taken from 15 sex- and age-matched healthy donors (HD).

### Human cell isolation

Peripheral blood mononuclear cells (PBMCs) were obtained from blood samples by the Lymphocytes Separation Media (cat. no. LSM-A; Capricorn, Germany) density gradient centrifugation method. Briefly, whole blood was diluted by the addition of an equal volume of PBS, blotted slowly onto separation medium, and centrifuged at 2000 rpm for 20 minutes at room temperature without stopping or acceleration. Cells were harvested from the interface, washed twice in phosphate buffered saline (PBS), then counted and prepared for staining. Tumor tissues and adjacent noncancerous tissues were dissociated into single cell suspensions using the human tumor dissociation kit (cat. no. 130-095-929; Miltenyi, Germany) according to the manufacturer’s instructions. Mediastinal lymph node tissues were mechanically dissociated into single cell suspensions. Red blood cells were removed from single cell suspensions using an ammonium chloride solution prepared in-house. The cells were filtered using a 70 μm cell strainer and then washed with PBS+ 2% fetal bovine serum (FBS) for subsequent flow cytometric assays.

### Mice

Wild-type (WT) C57BL/6J mice were provided by Medipol University Medical Research Center (Istanbul, Turkey). B6PL-Thy1.1 mice were obtained from The Jackson Laboratory (Bar Harbor, ME). Six-to eight-week-old C57BL/6J mice of both sexes were used as recipients of LLc1 tumor cells (tumor-bearing mice). B6PL-Thy1.1 mice of both sexes, six to eight weeks old, were used as donor mice for ILC2 isolation. For animal experiments, mice of the same age and sex were randomly assigned to experimental groups. Mice were housed under controlled conditions of temperatures of ∼18-23ºC with 40-60% humidity and a 12/12 h reverse light/dark cycle. Mouse care and experimental procedures were performed according to federal guidelines and protocols approved by Istanbul Medipol University Animal Experiments Local Ethics Committee (IMU-HADYEK). Tumor-bearing mice were monitored three times a week and at shorter intervals depending on the condition of the mice. Mice showing signs of stress, discomfort, pain, lethargy, inability to properly groom themselves, or inability to obtain food and/or water were immediately sacrificed.

### Tumor cell line

The murine Lewis lung carcinoma (LLc1) cell line was kindly provided by Prof. Dr. Güneş Esendağlı (Hacettepe University-Ankara/Turkey). The cells were maintained in complete medium consisting of RPMI 1640 medium (Sigma-Aldrich) supplemented with 2 mM L-glutamine (Sigma-Aldrich), 10% FBS and 1% penicillin–streptomycin (Gibco) at 37 °C in a humidified atmosphere with 5% CO_2_. Before injection, LLc1 cells (70–80% confluency) were harvested. Briefly, the cells were washed with PBS and detached using 0.25% trypsin-EDTA. Trypsin was neutralized with medium containing 10% FBS. After centrifugation, the cells were resuspended in PBS.

### Adoptive transfer of ILC2s

Briefly, for in vivo induction of ILC2s, donor mice (B6PL-Thy1.1) were hydrodynamically injected with 10 μg of a plasmid encoding murine IL-25 (pCMV3-mIL25, Sino Biological) as previously described (Frech et al. 2020). Mice were sacrificed three days post-injection, spleens and mesenteric lymph nodes (MLN) were harvested and mechanically dissociated into single cell suspensions. Red blood cells from spleen single cell suspensions were lysed using an ammonium chloride solution prepared in-house. MLN and spleen single cell suspensions were filtered through a 70μm cell strainer and washed with PBS+2% FBS. Lineage-positive cells were depleted from the resulting cell suspensions using the mouse direct lineage cell depletion kit (cat. 130-110-470, Miltenyi Biotec) according to the manufacturer’s instructions. The flow through containing lineage negative (Lin-) cells was washed with PBS+2% FBS before antibody staining and fluorescence-activated cell sorting (FACS) for CD45+ Lin-ICOS+ KLRG1+ ILC2s. FACS purified ILC2s were washed in PBS before injecting intravenously (i.v.) via the tail vein into tumor-bearing C57BL/6J recipient mice. Each injection consisted of 500,000 ILC2s in PBS. Each mouse received 3 injections per week for 4 weeks, and tissues were collected 5 days after the final injection for assessment of immune cell composition. For the survival experiment, mice continued to be monitored after the final ILC2 injection until natural death or euthanasia. Mice were euthanized when reaching humane endpoint (showing clinical signs of stress, discomfort, pain, lethargy, inability to properly groom themselves, inability to obtain food and/or water, hunching, weight loss above 20% of body weight, or a tumor volume above 1000mm^3^). For the experiment assessing the migration of ILC2s, each mouse received a single injection of ILC2s, and tissues were collected 24h later. All ILC2s for adoptive transfers were prepared fresh using the above protocol.

### Mouse Lymphodepletion

Tumor-bearing WT C57BL/6J mice were partially lymphodepleted by injecting cyclophosphamide (CTX, Sigma-Aldrich) intraperitoneally (i.p.) as a single dose at 200 mg/kg.

### Tumor volume assessment

6 -8-week-old WT C57BL/6J mice were inoculated subcutaneously (s.c.) into the axilla of the right forelimb with 1×10^5^ LLc1 cells in 100ul sterile PBS per mouse. Tumor growth was assessed three times per week by measuring tumor dimensions with calipers. Tumor volume was calculated using the following formula: V = (W^2^ × L)/2, W= Width; L= Length.

### Flow cytometry

For surface staining, both human and murine single cell suspensions were first stained with a viability marker (Fixable Viability Kit, Biolegend) according to manufacturer’s instructions to exclude dead cells, then Fc gamma receptors were blocked for 15 minutes with heat-inactivated human serum prepared in-house or anti-mouse CD16/32 (TruStain FcX, Biolegend) for human samples and mouse samples respectively, in PBS+2% FBS, and then incubated with the specific fluorochrome-labeled antibodies (all purchased from Biolegend unless otherwise noted) for 20 minutes at 4°C in the dark in PBS+2% FBS. For intracellular transcription factor staining, cells were fixed and permeabilized using a True-Nuclear Transcription Factor buffer kit (Biolegend) for 30 minutes at 4°C in the dark, washed twice with Perm buffer, and then stained for 30 minutes in Perm buffer. For intracellular cytokine staining on ILC2 cells, freshly sorted cells were stained ex vivo or after stimulation with carrier-free recombinant mouse IL-25 (Biolegend) at a concentration of 100 ng/ml in complete RPMI medium for 36 h. Both ex vivo and IL-25-stimulated cells were cultured in complete RPMI medium in the presence of phorbol-12-myristate-13-acetate (500 ng/ml) (Sigma-Aldrich) and ionomycin (250 ng/ml) (Sigma-Aldrich) for 4 h at 37°C immediately prior to staining. Brefeldin A (Biolegend) was added in the last 3 h of the culture. Cells were then stained surface stained as indicated above. Cells were then fixed, permeabilized and stained using True-Nuclear Transcription Factor buffer kit (Biolegend) according to manufacturer’s instructions. After staining, cells were washed with PBS+2% FBS, then resuspended in PBS+2% FBS for acquisition on a flow cytometer. Acquisition of human samples was performed on BD Influx. Acquisition of murine samples was performed on BD Symphony A1. Analyses of acquired data were performed using Flow Jo v10.9. Cell sorting was performed using BD Influx with >98% purity.

### Antibodies

For human samples, single-cell suspensions were stained with Zombie UV fixable viability kit (Biolegend), FITC-conjugated Hematopoietic Lineage cocktail prepared in-house (CD2 (TS1/8), CD3 (UCHT1), CD11b (ICRF44), CD14 (HCD14), CD15(HI98), CD16 (3G8), CD19 (HIB19), CD34 (561), CD56 (HCD56), CD123(6H6), CD20 (2H7), FcεRIα AER-37 (CRA-1)), PE-conjugated anti-CD127 (A019D5), APC/Cy7–conjugated anti-CD45 (2D1)), APC-conjugated anti-CD294 (CRTH2) (BM16), PE/Cy7-conjugated anti-HLA-DR (L243), BV421-conjugated anti-CD86 (BU63), BV510-conjugated anti-PD-L1 (29E.2A3), PE/Dazzle™ 594-conjugated anti-CD25 (BC96) and BUV661-conjugated anti-CD4 (SK3) or appropriate isotype controls. For intracellular staining, when sufficient numbers of cells were obtained, a separate staining was made with Zombie UV, BUV661-conjugated anti-CD4 (SK3) followed by intracellular staining with BV421-conjugated anti-FOXP3 (206D) or appropriate isotype controls.

For mouse experiments, the following antibodies were used for surface staining of ILC2 cells: FITC-conjugated Lineage cocktail prepared in-house (anti-CD3 (17A2), anti-CD5 (53-7.3), anti-B220 (RA3-6B2), anti-CD11b (M1/70), anti-CD11c (N418), anti-NK1.1 (PK136), anti-TER-119 (TER-119), anti-Gr1 (RB6-8C5), anti-CD170 (S17007L), anti-FcεRIα (Mar.01), anti-CD19 (6D5), anti-TCRβ (H57-597), anti-TCRγ/δ (UC7-13D5), PE-conjugated anti-CD45 (30.F11), APC-conjugated anti-KLRG (2F1/KLRG1), PE/Cy7 conjugated anti-CD278 (ICOS) (C398.4A), PerCP/Cyanine5.5-conjugated anti-CD90.1 (Thy1.1) (OX-7), Zombie UV fixable viability dye. For intracellular staining of ILC2 cells, the following antibodies were used: PE-conjugated anti-IL-5 (TRFK5), APC/Cy7-conjugated anti-IL-13 (eBioscience, clone eBio13A), PE/Dazzle 594-conjugated anti-IL-4 (11B11), and PE/Cy7-conjugated anti-GATA3 (eBioscience, clone TWAJ) or appropriate isotype controls. For immune cell composition of tissues obtained from tumor-bearing mice, the following antibodies were used for surface staining: PE/Cy7-conjugated anti-CD3 (eBioscience, clone 145-2C11), BV785-conjugated anti-CD4 (GK1.5), BV510-conjugated anti-CD8 (GK1.5), AF594-conjugated anti-CD11b (M1/70), FITC-conjugated anti-CD11c (N418), PE-conjugated anti-Ly6C (HK1.4), APC-conjugated anti-Ly6G (1A8), PE/Cy5-conjugated anti-MHCII (M5/114.15.2), Zombie UV fixable viability dye. For intracellular staining, AF700-conjugated anti-FOXP3 (MF-14) was used.

### Murine cell isolation

A maximum of 1g of tumor and lung tissue was rinsed with PBS, cut into small pieces using scissors and placed in a gentleMACS C tube (Miltenyi Biotec) in 5 ml RPMI1640 containing 100 μl 10X collagenase/hyaluronidase enzyme mixture (cat. 07912, Stemcell). The C tubes were then placed on a gentleMACS dissociator (Miltenyi Biotec) and the appropriate programs were run according to the mouse tumor dissociation kit protocol and mouse lung dissociation kit protocol (both from Miltenyi Biotec) for tumors and lungs respectively. The samples were then incubated for 45 min at 37°C while rotating. The digested fragments were manually dissociated further by mashing them with a syringe plunger onto a 70 μm cell strainer and the cells were washed in PBS+ 2% FBS. Erythrocytes were removed with an ammonium chloride solution prepared in-house. Mononuclear leukocytes from tumors were isolated by performing a Histopaque 1083 (Sigma-Aldrich) density gradient centrifugation according to the manufacturer’s instructions. Obtained single cell suspensions were then washed in cold PBS+ 2% FBS buffer before staining. Liver tissue was dissociated by following the ‘preparation of single cell suspensions from mouse liver’ protocol from Miltenyi Biotec. Cells were then washed in cold PBS+ 2% FBS buffer before staining. Spleen and lymph nodes were mechanically dissociated then filtered through a 70-μm cell strainer. Red blood cells from the spleen were removed using ammonium chloride solution prepared in-house. Samples were washed in cold PBS+ 2% FBS buffer prior to staining.

### Statistical analysis

Statistical analyses and representations were performed using Prism (GraphPad Version 9.0) and are detailed in the figure legends.

## RESULTS

### ILC2 cells and Foxp3^+^ Tregs are enriched in NSCLC patients

Total helper ILCs were identified as live CD45^+^ Lin^-^ CD127^+^ cells, among which ILC2 cells were identified as CRTH2^+^ cells, in line with the literature and previous studies (Carrega et al. 2015; Shen et al. 2021; Vivier et al. 2018). Representative gating strategies for total ILC helpers and ILC2 cells are shown in Figure 1A. Total ILC helper frequencies among live CD45^+^ cells were similar between tissues obtained from NSCLC patients (peripheral blood (PB), lung-draining mediastinal lymph node (LN), tumor tissue and adjacent normal lung tissue) and PB obtained from HDs (Figure 1B). Within NSCLC patient samples, the proportion of ILC helper cells among live CD45^+^ cells was significantly higher in normal lung tissue as compared to lung-draining mediastinal LNs and PB, however there was no significant difference between their proportions in normal lung tissue and tumor tissue, tumor tissue and LN, nor tumor tissue and PB (Figure 1B). The aforementioned results are consistent with the recent study by Shen at al. (Shen et al. 2021). Next, we analyzed the frequency of ILC2 cells among total ILC helpers. ILC2 cells were found to be enriched in all tissues obtained from NSCLC patients (except LN), as compared to HDs (Figure 1C), in line with the previous report by Shen et al. In NSCLC patients, ILC2 cell frequencies were found to be significantly higher in the lung (both normal tissue and tumor tissue) and PB as opposed to the mediastinal LNs (Figure 1C), consistent with previous reports on their tissue distribution (Simoni et al. 2017). As opposed to a previous study by Carrega et al. (Carrega et al. 2015), there was no significant difference between the frequencies of ILC2 cells in the tumor tissue as compared to matched normal lung tissue (Figure 1C). This discrepancy may be due to the difference in the number of samples analyzed in the study by Carrega et al. (n=16) as compared to our study (n=28). While the percentages of ILC2 cells among ILC helpers were similar between tumor tissue and matched normal lung tissue obtained from NSCLC patients, their absolute numbers per mg of tissue were significantly higher in tumor tissue as compared to adjacent normal lung (Figure 1D), showing a preferential accumulation of ILC2 cells within tumors.

**FIGURE 1.**
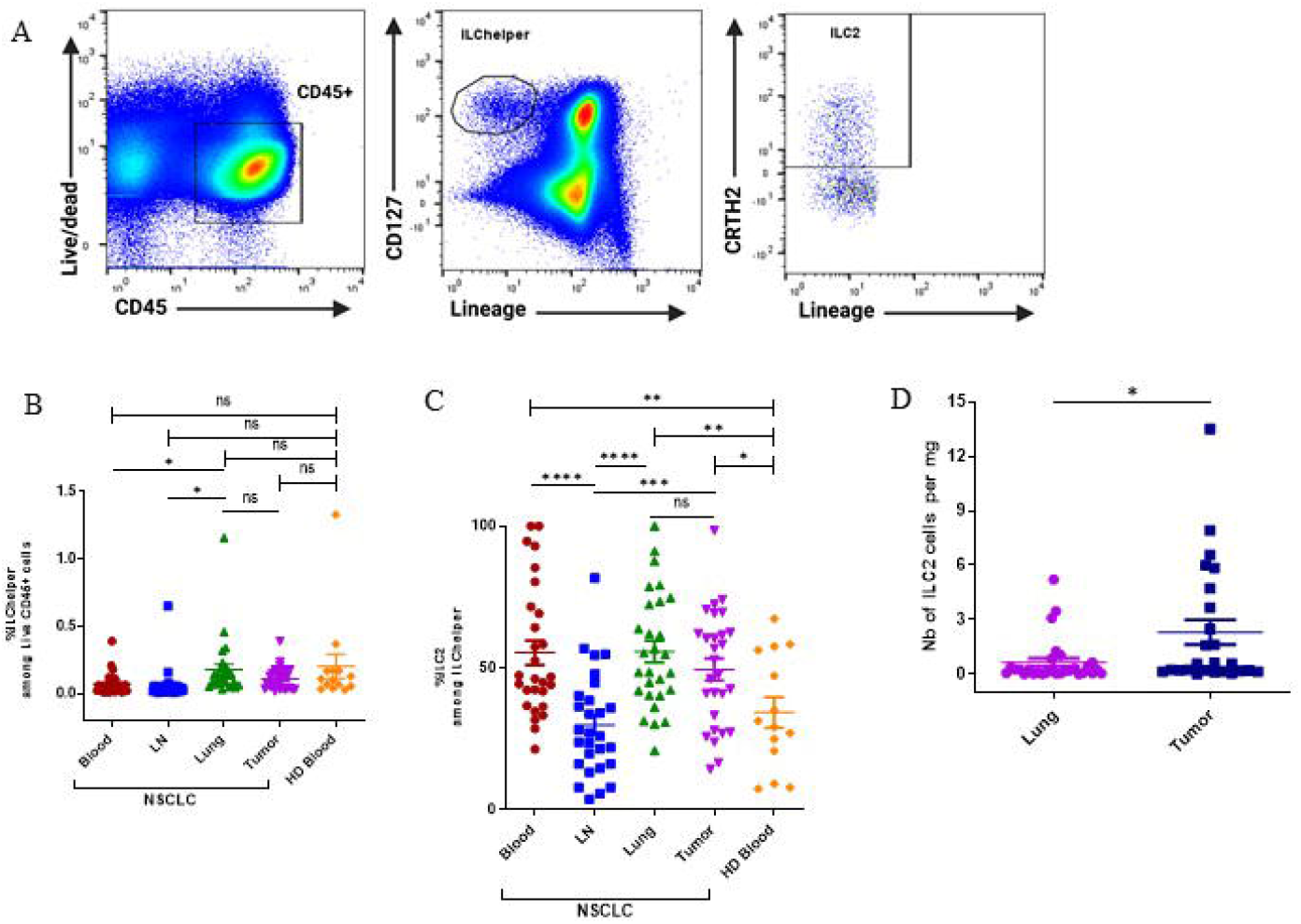
Distribution of group 2 innate lymphoid cells in human NSCLC tissues and healthy donors. (A) Representative gating strategies for total ILC helpers (live CD45^+^ Lin (CD2, CD3, CD11b, CD14, CD15, CD16, CD19, CD34, CD56, CD123, CD20, FcεRIα.)^−^ CD127^+^ and ILC2 cells (live CD45^+^ Lin^−^ CD127^+^ CRTH2^+^). (B) Frequencies of total ILC helpers among CD45^+^ cells in tissues obtained from NSCLC patients (for each group, n=28) and peripheral blood obtained from HD (n=l4). (C) Percentages of ILC2s among total ILC helpers in NSCLC tissues (for each group, n=28) and HD (n=14). (D) Absolute numbers of ILC2 cells per mg of tissue in tumor tissue and matched normal lung tissue obtained from NSCLC patients (for each group, n=28). LN: tumor-draining mediastinal lymph node; Lung: normal lung tissue adjacent to tumor tissue; HD: healthy donor. ns, not significant; *:p<0.05; **:p<0.01; ***:p<0.001; ****p<0.0001. P-values were obtained in an unpaired two-tailed Student’s t-test. Horizontal bars mark mean. Error bars mark sem.

Murine ILC2 cells have been reported to express MHC Class II and to be able to function as antigen-presenting cells (APCs) (Hepworth et al. 2013; Mirchandani et al. 2014; Oliphant et al. 2014; Symowski and Voehringer 2019). In humans, only one study reported MHC Class II and CD80 expression on ILC2 cells as being elevated in patients with acute exacerbation of chronic pulmonary obstructive pulmonary disease (AECOPD) (Jiang et al. 2019). Moreover, ILC2 cells stimulated by IL-33 were found to upregulate PD-L1 (Taylor et al. 2017). PD-L1 is a ligand for PD-1, which is a known checkpoint inhibitor of CD4^+^ T cells that plays important roles in diminishing anti-tumor immunity. We therefore wanted to assess whether ILC2 cells in NSCLC patients may have immunosuppressive capacities by interacting directly with CD4^+^ T cells via MHC II, CD86 and PD-L1. To this end, we assessed the expression of HLA-DR, CD86 and PD-L1 on ILC2 cells in the various tissues obtained from NSCLC patients as well as in HDs (Suppl. Fig. 1). As opposed to murine ILC2 cells and ILC2 cells in AECOPD, ILC2 cells in PB of HDs and tumors of NSCLC patients did not express HLA-DR, CD86 nor PD-L1 (Suppl. Fig. 1). In NSCLC patients, the lack of expression of HLA-DR, CD86 nor PD-L1 by ILC2 cells was irrespective of the tissue of origin (data not shown). However, ILC2 cells from NSCLC and HD samples did express CD25 (Suppl. Fig 1), consistent with the literature (Spits et al. 2013). CD25 expression by ILC2 cells in samples obtained from NSCLC patients was irrespective of the tissue of origin (data not shown).

Next, we assessed the frequencies of CD4^+^ T lymphocytes among live lymphocytes and frequencies of Foxp3^+^ regulatory T cells among total CD4^+^ T cells (Fig. 2). While the proportions of CD4^+^ T lymphocytes were reduced in all tissues except LNs (where the proportions were similar to HDs) obtained from NSCLC patients as compared to HDs (Fig. 2A), Foxp3^+^ regulatory T cells were found to be enriched in all tissues except PB (where the proportions were similar to HDs) obtained from NSCLC patients as compared to HDs (Fig. 2B). These results are consistent with an accumulation of immune suppressor cells concomitantly with a reduction in effector immune cells preferentially within NSCLC patients.

**FIGURE 2.**
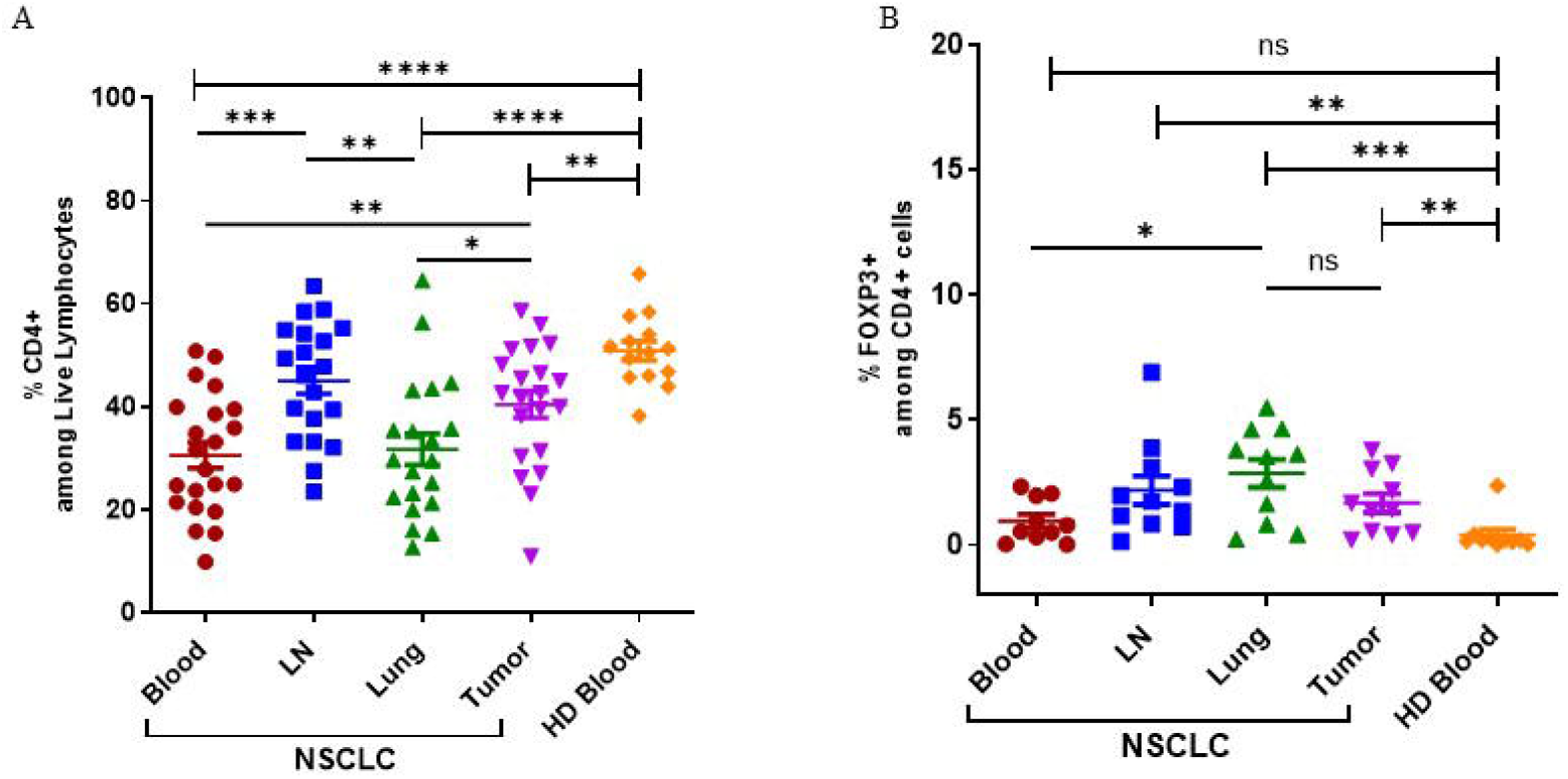
Frequencies of CD4^+^ T cells and Foxp3^+^ regulatory T cells in NSCLC patients and HD. (A) Frequency of CD4^+^T cells in tissues obtained from NSCLC patients (peripheral blood (PB), n=22; LN, n=20; Lung; n=20; Tumor, n=21) and PB obtained from HD (n=l4). (B) Frequency of Foxp3^**+**^ regulatory T cells among CD4^**+**^T lymphocytes in tissues obtained from NSCLC patients (PB, n=l0; LN, n=11; Lung; n=11; Tumor, n=11) and PB obtained from HD (n=10). LN: tumor-draining mediastinal lymph node; Lung: normal lung tissue adjacent to tumor tissue; HD: healthy donor. Horizontal bars mark mean. Error bars mark sem. P-values were calculated with unpaired two tailed Student’s t-test. *p < 0.05; **p < 0.01; ***p< 0.001; ****p< 0.0001.

Taken together, our findings show that ILC2 cells as well as Foxp3^+^ Tregs are preferentially enriched in NSCLC patients, especially tumors, as compared to HDs, suggesting that ILC2 cells may bear a tumor-promoting function in NSCLC.

### Adoptive transfer of ILC2 cells in tumor-bearing mice promotes tumor growth and reduces survival

In order to test our hypothesis that ILC2 cells bear tumor-promoting properties, we decided to examine the effect of adoptive transfer of ILC2 cells in lung tumor-bearing mice using the well-established Lewis lung carcinoma (LLc1) heterotopic model of lung cancer. In order to obtain sufficient numbers of ILC2 cells for adoptive transfer, endogenous ILC2 cells were expanded *in vivo* by hydrodynamic gene delivery (hgd) of an IL-25-encoding plasmid (pCMV-IL25) as previously described (Frech et al. 2020). Such ILC2 cells were identified as CD45^+^ Lin^-^ KLRG1^+^ ICOS^+^ cells and were recovered from mesenteric LNs (MLNs) and spleen as previously described (Frech et al. 2020). A representative example of the *in vivo* expansion and phenotypic profile of such ILC2 cells following hgd is shown in Fig. 3A. These cells were high producers of IL-4 and IL-13, but not IL-5 (Fig. 3B and C). Their cytokine production profile remained the same when cells were restimulated with murine IL-25 *in vitro* (Fig. 3B and C). Moreover, these cells expressed the master transcription factor GATA3 (Figure 3D). Altogether, these results show that CD45^+^ Lin^-^ KLRG1^+^ ICOS^+^ cells obtained after hgd with IL-25-encoding plasmid were true ILC2 cells that expressed type 2 cytokines and GATA3. Moreover, their expansion by IL-25 and high KLRG1 expression reminisce of the inflammatory subset of ILC2 cells (iILC2) previously described (Y. Huang and Paul 2016).

**FIGURE 3.**
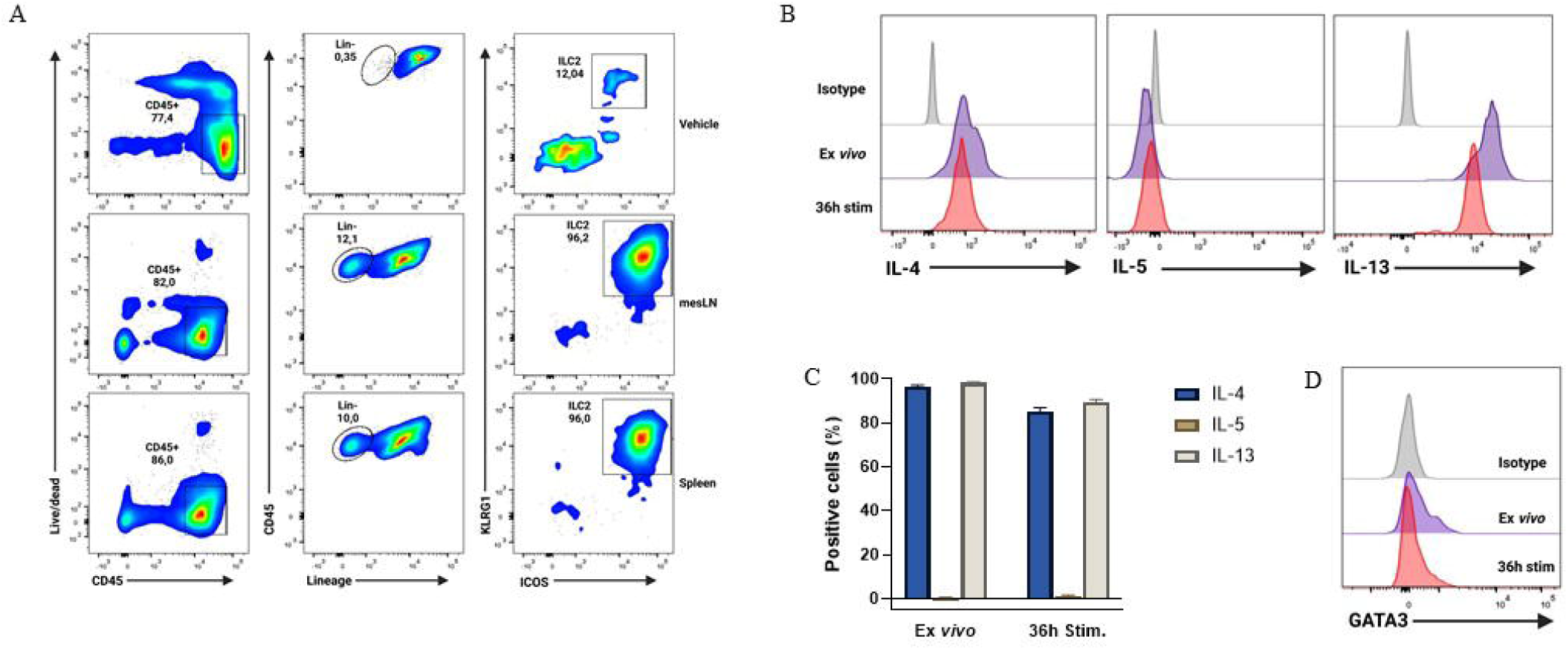
Phenotype and cytokine profile of in vivo expanded ILC2s after hgd with pCMV-IL25 in mice. (A) Gating strategy for the identification of ILC2 cells (live CD45^+^ Lin (CD3, CD5, B220, CD11b, CD11c, NKl.1, TER-119, Gr1, CD170, FcεRIα, CD19, TCRβ TCRγ/δ)-KLRG1^+^ ICOS^+^) 3 days after hgd with vehicle (top panel) or pCMV-IL25 (bottom 2 panels) in B6PL-Thyl.l mice. mesLN: mesenteric lymph node. (B) Intracellular cytokine analysis of in vivo expanded ILC2 cells from mesLN, stimulated with PMA/ionomycin for 4h directly after obtention of mononuclear cell suspension (ex vivo, purple line) or after 36h restimulation with murine recombinant IL-25 (36h stim, red line). Results are shown as gated on live Lin-ICOS^+^ cells. Appropriate isotype controls (gray lines) are shown. One representative example out of 3 is shown. (C) Pooled data (n=3) are shown. Bars represent the mean values of the percentages of positive cells ± sem. (D) Representative example of GATA3 transcription factor expression by in vivo expanded ILC2 cells stained directly after obtention of mononuclear cell suspension (ex vivo, purple line) or after 36h restimulation with murine recombinant IL-25 (36h stim, red line) as stated in B. Appropriate isotype (gray line) is used as control. One representative example out of 3 is shown.

In order to temporarily free up niches for adoptively transferred cells, we decided to partially lymphodeplete recipient mice by administering cyclophosphamide (CTX) i.p. one day prior to adoptive transfer. First, we wanted to assess the migration potential of ILC2 cells adoptively transferred i.v. into tumor-bearing mice. To this end, we transferred Thy1.1^+^ ILC2 cells into tumor-bearing Thy1.2^+^ recipient mice one day after CTX or PBS vehicle injection (Figure 4A). The recipient mice were sacrificed 24h after ILC2 transfer and various organs were collected for staining with anti-Thy1.1 antibody (Figure 4A). As shown in Figure 4B, transferred ILC2 cells migrated to all tissues of recipient mice, however the frequencies of transferred cells were highest in tumor tissue and spleen. In mice that were not lymphodepleted prior to ILC2 adoptive transfer, the percentage of transferred cells was overall very low, but the same pattern of highest migration to tumor and spleen was observed (Figure 4B, lower panels). Lymphodepletion with CTX prior to adoptive transfer enabled a 3-fold increase in the frequency of transferred cells that could be recovered in tumors (Figure 4B, 0.083% in vehicle-treated mice vs. 0.28% in CTX-treated mice).

**FIGURE 4.**
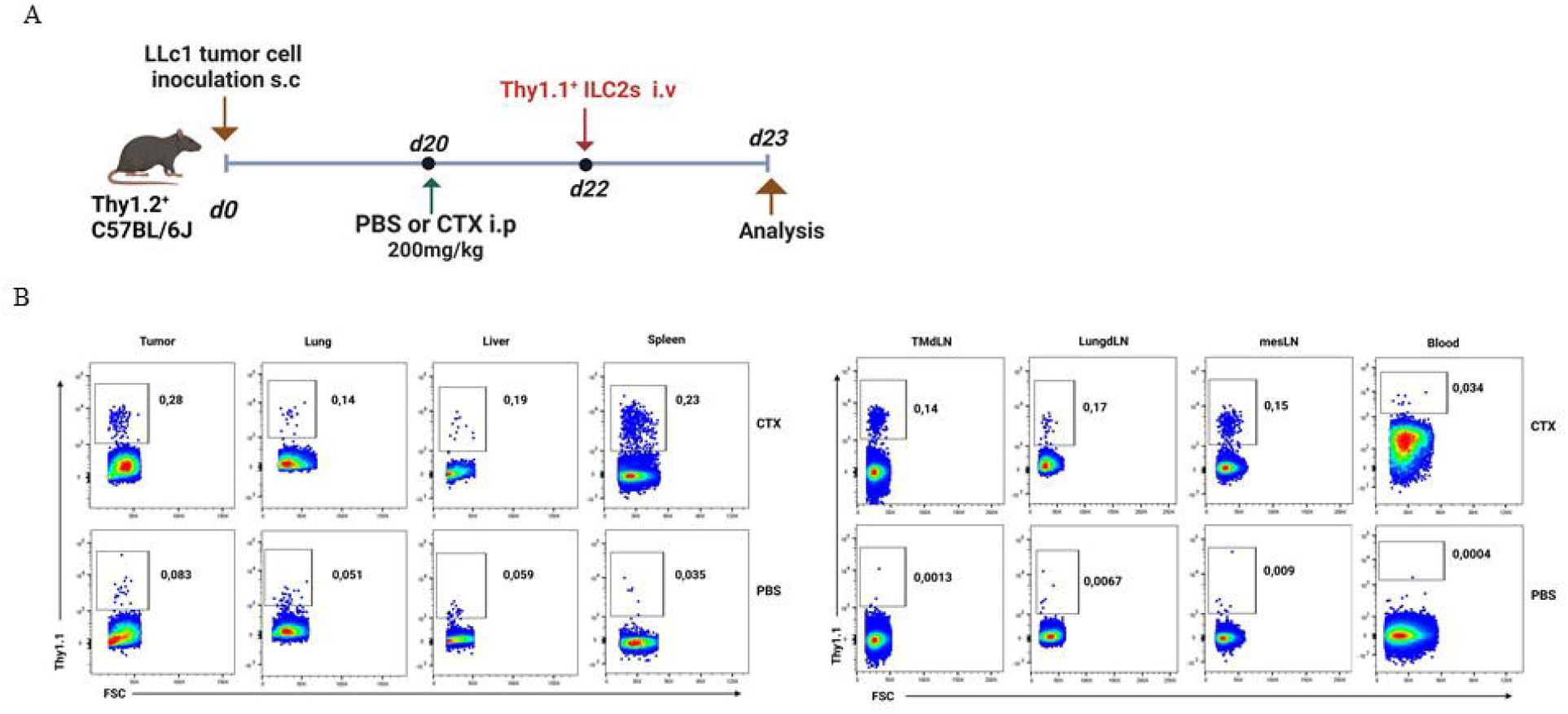
Migration of adoptively-transferred ILC2 cells in tumor-bearing mice. **(A)** Schematic representation of experimental design. Tumor-bearing Thyl.2^+^C57BL/6J mice were partially lymphodepleted by receiving CTX at 200mg/kg in PBS i.p. Controls received PBS only. 48h later, mice were adoptively transferred with 5×10^5^ Thyl.1^+^ ILC2 cells i.v. in the tail vein. Recipient mice were sacrificed 24h after adoptive transfer and tissues were collected. i.p: intraperitoneal; i.v.: intravenous. (B) Representative flow cytometry plots showing the percentage of adoptively-transferred Thy1.1^+^ ILC2s in various tissues obtained from the lymphodepleted (CTX, upper panels, n=3) Thyl.2^+^ tumor-bearing recipient mice and non-lymphodepleted (PBS, lower panels, n=3) recipient mice. Plots are gated on live lymphocytes.

Next, in order to assess the effect of ILC2 cells on tumor growth and survival, we transferred freshly sorted ILC2 cells or vehicle into lymphodepleted tumor-bearing mice twice per week for 4 weeks (Figure 5A). The purity of ILC2 cells after cell sorting was at or above 98% (Suppl. Fig. 2). Adoptive transfer of ILC2 cells had no effect on tumor growth in mice that did not receive CTX treatment (Suppl. Fig. 3). This is probably due to the fact that only very low numbers of ILC2 cells could reach the tumor in non-lymphodepleted tumor-bearing mice (Figure 3B, lower panels). Although initially the tumor volume was similar in mice that received ILC2 cells and controls, it started to become significantly higher in mice that received ILC2 cells as compared to those that received vehicle after the second week of treatment (Figure 5B). Overall, ILC2 adoptive transfer led to significant higher tumor burden (Figure 5B). Moreover, the survival of mice was significantly reduced when ILC2 cells were transferred (Figure 5C).

**FIGURE 5.**
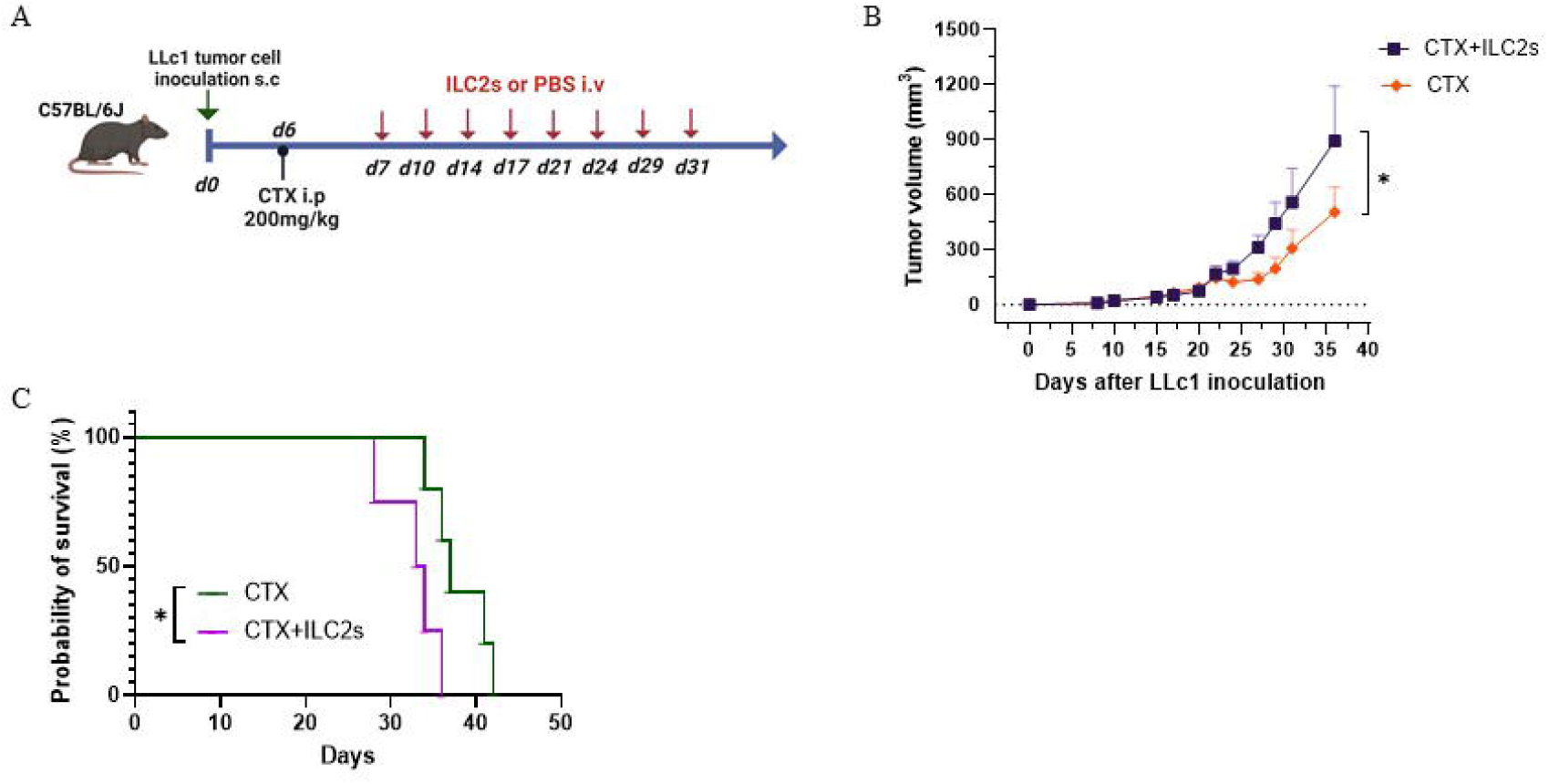
Adoptive transfer of ILC2 cells increases tumor burden and decreases survival. (A) Schematic representation of experimental design. 6-8-week old C56BL/6J mice were inoculated with l×10^5^ LLc1 lung tumor cells on day 0, then partially lymphodepleted with CTX on day 6. Starting one day after CTX administration, mice received 5×10^5^ freshly sorted ILC2 cells or vehicle 2 times per week for 4 weeks. s.c:subcutaneous; i.p: intraperitoneal; i.v.: intravenous. (B) Tumor volume in mice that received ILC2s (CTX+ILC2s; n=9) vs. mice that received vehicle (CTX; n=10). Data show mean ±sem. Data were pooled from 3 independent experiments with ≥3 mice per group. (C) Survival of mice mice that received ILC2s (CTX+ILC2s; n=4) vs. mice that received vehicle (CTX; n=5).*:p<0.05. p values were determined by Student’s I-test (B) and log-rank test (C).

### Adoptive transfer of ILC2 cells leads to accumulation of suppressive M-MDSCs into tumors

Next, we sought to determine the TME composition after ILC2 transfer in CTX-lymphodepleted tumor-bearing mice. To this end, we transferred freshly sorted ILC2 cells or vehicle into lymphodepleted tumor-bearing mice twice per week for 4 weeks. Recipient mice were sacrificed 5 days after the last ILC2 injection and tumors, spleens and lymph nodes (LNs) were collected for analysis (Figure 6A). As shown in Figure 6B, the percentages of CD4^+^, CD8^+^ and Foxp3^+^ Treg T lymphocyte subsets were similar in tumors of mice that received ILC2 cells or PBS. Figure 6C shows the percentages of myeloid cells in tumors. ILC2 cell transfer did not alter the frequencies of conventional dendritic cells (cDCs) nor granulocytic MDSCs (G-MDSCs), however there was a significant increase in the frequencies of monocytic MDSCs (M-MDSCs) in mice that received ILC2 cells as compared to controls (Figure 6C). This is in line with a previous report on IL-25-activated ILC2 cells promoting colorectal cancer by activating M-MDSCs (Jou et al. 2022). The accumulation of M-MDSCs in tumor-bearing mice that received ILC2 cells was not systemic but rather specific to tumors since there was no significant difference in their frequencies within spleens and LNs (Suppl. Fig. 4 and data not shown). This result is consistent with the higher migration potential of ILC2 cells into tumors as compared to other tissues (Figure 4B).

**FIGURE 6.**
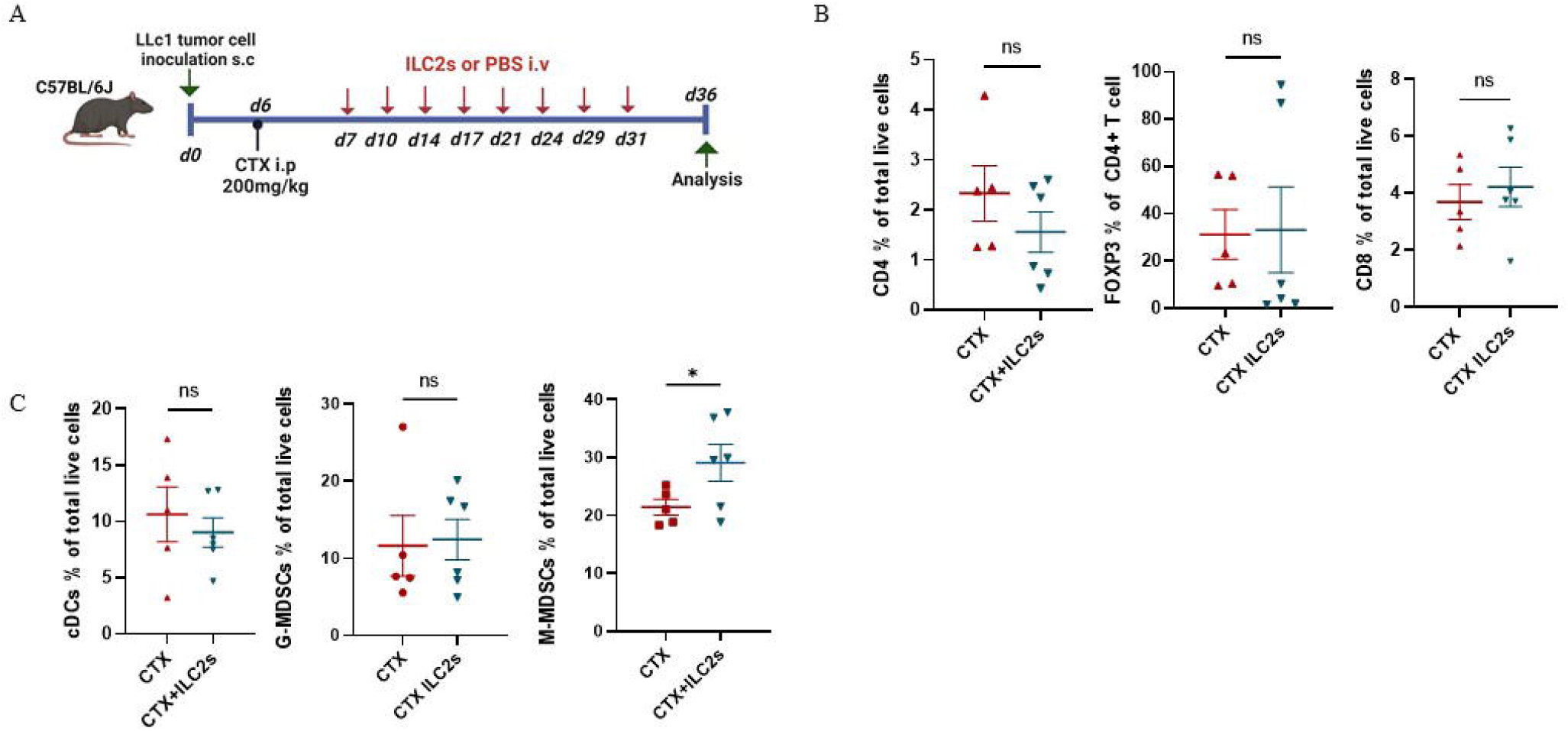
Adoptive transfer of ILC2 cells promotes the accumulation of M-MDSCs in tumors. (A) Schematic representation of experimental design. 6-8-week old C56BL/6J mice were inoculated with l×10^5^ LLc1 lung tumor cells on day 0, then partially lymphodepleted with CTX on day 6. Starting one day after CTX administration, mice received 5×10^5^ freshly sorted ILC2 cells or vehicle 2 times per week for 4 weeks. Mice were sacrificed 5 days after the last injection and tumors were collected. s.c: subcutaneous; i.p: intraperitoneal; i.v.: intravenous. (B) Frequency of tumor-infiltrating CD4 T cells identified as CD3^+^ CD4^+^, CD8 T cells identified as CD3^+^ CD4^+^ and CD4 regulatory T (Treg) cells identified as CD3^+^ CD4^+^ Foxp3^+^ in vehicle (CTX, n=5)- and ILC2-treated (CTX+ILC2, n=6) tumor-bearing mice. (C) Frequency of tumor-infiltrating conventional DCs (cDCs) identified as CD11c^+^ MHCII^+^ cells, granulocytic myeloid derived suppressor cells (G-MDSCs) identified as CD3^−^ CD11b^+^ Ly6G^+^ Ly6C^low^, and monocytic MDSCs (M-MDSCs) identified as CD3^−^ CD11b^+^ Ly6G^−^ Ly6C^hi^ in vehicle (CTX, n=5)- and ILC2-treated (CTX+ILC2, n=6) tumor-bearing mice. Horizontal bars mark mean. Error bars mark sem. Statistical significance was calculated by unpaired two-tailed t-test. ns, not significant; *:p<0.05.

Overall, our results show that IL-25-activated ILC2 cells preferentially migrate to tumors, and that they promote tumor growth and reduce survival of tumor-bearing mice. Moreover, we found that M-MDSCs accumulated into tumors following adoptive transfer with ILC2 cells, suggesting that ILC2s may actively recruit M-MDSCs into tumors. This may be due to the high production of IL-4 and IL-13 by ILC2 cells (Fig. 3B and C) as it has previously shown that ILC2s mediate the recruitment of MDSCs via IL-13 (Chevalier et al. 2017; Trabanelli et al. 2017) and that they induce the expansion and activation of M-MDSCs via the activation of STAT6 through the binding of the IL4Ralpha (Gabrilovich and Nagaraj 2009; Jou et al. 2022).

## DISCUSSION

Here, we found that ILC2 cells accumulate in lung tumors and that they promote lung tumor growth, in humans and mice respectively. In humans, we assessed the frequencies of ILC2 cells in tumor tissue, adjacent normal lung tissue, mediastinal LN and PB of NSCLC patients, as well as in PB from HDs. This is the first time that such a comprehensive study has been done. We found that ILC2 cells were enriched in tumors and PB of NSCLC patients as compared to PB from HDs, confirming previous results by Shen et al. (Shen et al. 2021) using a different cohort of patients. In addition to the results provided by Shen et al., we assessed the frequencies of ILC2 cells in lung-draining mediastinal LNs and normal lung tissue adjacent to tumor of NSCLC patients, and found that ILC2 cells were enriched in all tissues of NSCLC patients as compared to HDs except for LNs. This is consistent with previous reports on the localization of ILC2 cells in human tissues (Simoni et al. 2017). We found that there was no significant difference between the frequencies of ILC2s in tumor tissue and adjacent lung tissue. This result is in contrast with a previous report by Carrega et al. (Carrega et al. 2015), and may be explained by the nearly double number of samples assessed in our study. Moreover, we report absolute numbers of ILC2s per mg of tissue for the first time and found that they were significantly higher in tumors as compared to normal lung tissue. Additionally, we assessed the frequencies of CD4^+^ T cells and Foxp3^+^ Tregs in the obtained NSCLC and HD samples. We found that total CD4^+^ T cells were reduced while Foxp3^+^ Tregs were enriched in NSCLC patients, also especially in tumors, as compared to HDs. To the best of our knowledge, this is the first comprehensive report concomitantly comparing Foxp3^+^ Treg and ILC2 frequencies in various tissues of NSCLC patients and HDs by flow cytometry.

While the role of ILC2 cells in allergy and parasitic infections is now well-established, their role in cancer is still controversial (Magnusson and Bahhar 2023). In a murine model of heterotopic lung cancer expressing IL-33, ILC2-deficient mice, which were generated by reconstituting their bone marrow with RORalpha^-/-^ hematopoietic stem cells, showed increased tumor growth and metastasis, suggesting an anti-tumor role for IL-33-activated ILC2 cells (Saranchova et al. 2018). In other cancer types such as pancreatic cancer, melanoma, lymphoma or colon cancer, IL-33-activated ILC2 cells were reported to bear protective functions (reviewed in Magnusson and Bahhar 2023 and Trabanelli et al. 2019). However, recent evidence suggests an opposite function for IL-25-activated ILC2 cells. In a mouse model of colorectal cancer, IL-25 activation of ILC2 cells promoted intestinal tumorigenesis by recruiting and activating immunosuppressive M-MDSCs (Jou et al. 2022). Here, the discrepancy may be explained by three different reasons: first, different cancer types have disparate TMEs, immune compositions and immune reactions. Second, different models of the same cancer type have distinct genetic mutations that can influence the immune environment and therefore the immune reactions to tumors. Finally, differing activation signals received by ILC2 cells may lead to different outcomes. It has recently been proposed that IL-33-responsive ILC2 cells represent ‘natural ILC2s’ (nILC2s) while IL-25-responsive ILC2 cells constitute ‘inflammatory ILC2s’ (iILC2s), both with distinct tissue localization, cell surface marker expression, and potentially distinct functions (Y. Huang and Paul 2016). The presence of different subsets of ILC2 cells in IL-33-expressing and IL-25-expressing cancers could explain the discrepancies in their observed functions.

The ILC2 cells used in our study are consistent with the described iILC2s by Huang et al (Y. Huang and Paul 2016). Indeed, our ILC2 cells are induced by IL-25, they express high levels of KLRG1, they can be found in the mesenteric LN (MLN) and spleen, and they produce large amounts of IL-13 and IL-4. Our ILC2s preferentially migrated to tumors and promoted tumor growth as well as reduced survival in lung tumor-bearing mice, possibly by recruiting M-MDSCs into tumors, consistent with the findings by Jou et al. (Jou et al. 2022) in CRC. Thus, iILC2 (or IL-25-activated ILC2 cells) may bear tumor-promoting functions while nILC2 (or IL-33-activated ILC2 cells) cells may have opposite functions in lung cancer. Further studies are needed to better understand the roles of iILC2s vs nILC2s in cancer.

The present study has some limitations. We found that, in mice, M-MDSCs accumulated into tumors following adoptive transfer with ILC2 cells, suggesting that ILC2 cells may recruit M-MDSCs. Several reports have shown that IL-13 produced by ILC2 cells can mediate the selective recruitment of M-MDSCs that express IL-13Ralpha1 (Chevalier et al. 2017; Trabanelli et al. 2017). The ILC2 cells used in our study produce large amounts of IL-13, however we did not show whether the recruitment of M-MDSCs into tumors after ILC2 adoptive transfer was dependent on IL-13 production by ILC2 cells. Further well-designed studies are therefore needed to test for this hypothesis. Moreover, the ILC2 cells used in our study also produce large amounts of IL-4. Jou et al. (Jou et al. 2022) showed that ILC2-derived IL-4 and IL-13 promoted the suppressive capacities of M-MDSCs by increasing arginase 1 (Arg1) expression, therefore the ILC2 cells used in our study may not only promote the recruitment of M-MDSCs but they may also increase their immunosuppressive capacities. Additional studies, beyond the scope of the present contribution, are needed to better understand the direct effects of ILC2 cells on M-MDSCs. Finally, although we found that ILC2 cells and Foxp3^+^ Tregs were enriched in NSCLC patients, we did not assess the frequencies of M-MDSCs in these patients. Nevertheless, previous studies did report higher frequencies of M-MDSCs both in the peripheral blood and in tumors of NSCLC patients as compared to HDs (Yamauchi et al. 2018; Zadian et al. 2021) as well as an association between increased ILC2s and MDSCs in patients with lung cancer (Wu et al. 2017).

In humans, whether there are two subsets of ILC2 cells, namely nILC2 and iILC2 such as in mice, is unknown. Indeed, human ILC2 cells are defined by the expression of CD45, CRTH2 and the lack of expression of lineage markers (Vivier et al. 2018). They also express CD25, ICOS, ST2 (a subunit of IL-33R) and IL-17RB (a subunit of IL-25R) (Vivier et al. 2018). While IL-25 and IL-33 have been shown to induce distinct activation profiles in human ILC2 cells *in vitro* (Camelo et al. 2017), whether IL-25-responsive ILC2s and IL-33-responsive ILC2s constitute different subsets of ILC2 cells in humans is unknown. In NSCLC patients, IL-25R and ST2 expression by ILC2 cells has never been assessed. Although we report CD25 expression in ILC2 cells from NSCLC patients, we did not assess IL-25R nor ST2 expression in our study.

Furthermore, due to the rarity of ILC2 cells in clinical samples, we could not assess whether ILC2 cells from NSCLC patients were more responsive to IL-25 or IL-33. At the steady-state, murine lungs are known to bear nILC2 cells that are responsive to IL-33, however, under inflammatory conditions, IL-25-responsive iILC2 cells are quickly induced (Y. Huang and Paul 2016). Whether this is the case in humans as well, and especially in NSCLC patients, remains to be determined.

In summary, our data supports a tumor-promoting role for ILC2 cells both in humans and in mice. In mice, IL-25-responsive ILC2 cells were able to migrate to tumors, promote tumor growth, reduce the survival of mice, and they were associated with an accumulation of M-MDSCs into tumors, suggesting that they may actively mediate their recruitment. In humans, ILC2 cells were found to be enriched in NSCLC patients, especially within tumors. Furthermore, concomitant with ILC2 cells, the frequency of immunosuppressive Foxp3^+^ Tregs was also increased in NSCLC patients as compared to HDs. Overall, our results suggest that ILC2 cells may bear pro-tumoral functions by recruiting immune-suppressive cells, and that targeting ILC2 cells might represent a novel immunotherapy for patients with NSCLC.

## DATA AVAILABILITY STATEMENT

The raw data supporting the conclusions of this article will be made available by the authors, without undue reservation.

## AUTHOR CONTRIBUTIONS

Conception, design and supervision: FCM. Development of methodology: IB. Acquisition of human material: AT, MZG, AÇ, IB. Acquisition of murine data: IB, ZE, OK. Analysis and interpretation of data: IB, FCM. Writing, review and/or revision of the manuscript: FCM, DD. All authors contributed to the article and approved the submitted version.

## FUNDING

This work was supported by two grants awarded by the Scientific and Technological Research council of Turkey (TÜBITAK) with grant numbers 119S136 and 217S770.

## ACKNOWLEDGMENTS

We thank Emre Vatandaşlar for assistance with cell sorting and the whole staff and students of REMER center at Istanbul Medipol University for their invaluable help in this study.

## SUPPLEMENTARY MATERIAL

**Supplementary Table S1.** Clinicopathological characteristics of patients with NSCLC.

**Supplementary Figure 1.** Phenotypic characterization of ILC2 cells in NSCLC patients and HD. HLA-DR, CD86, PD-L1, and CD25 expression by ILC2 cells determined by flow cytometry analysis of cell surface marker gated on live CD45^+^ Lin^-^ CD127^+^ CRTH2^+^. Specific antibodies (purple and red lines) or appropriate isotype controls (gray lines) are shown. One representative example out of n=22 (NSCLC) and n=15 (HD) is shown. NSCLC: tumor sample from non-small cell lung cancer patient; HD: peripheral blood sample from healthy donor.

**Supplementary Figure 2.** Cell sorting for in vivo expanded ILC2 cells 3 days after hgd with pCMV-IL25 in mice. (A) Pre-sort purity of ILC2 cells (live CD45^+^ Lin (CD3, CD5, B220, CD11b, CD11c, NK1.1, TER-119, Gr1, CD170, FcεRIα, CD19, TCRβ, TCRγ/δ)^-^ KLRG1^+^ICOS^+^) in pooled MLN and spleen cell suspensions that were depleted of lineage positive cells with the mouse direct lineage cell depletion kit. One representative example out of n>20 is shown. (B) Post-sort purity of ILC2 cells (live CD45^+^ Lin (CD3, CD5, B220, CD11b, CD11c, NK1.1, TER-119, Gr1, CD170, FcεRIα, CD19, TCRβ, TCRγ/δ)^-^ KLRG1^+^ ICOS^+^) in pooled MLN and spleen cell suspensions. One representative example out of n>20 is shown.

**Supplementary Figure 3.** Adoptive transfer of ILC2 cells has no effect on tumor growth in non-lymphodepleted recipient mice. (A) Schematic representation of experimental design. 6-8-week old C56BL/6J mice were inoculated with 1×10^5^ LLc1 lung tumor cells on day 0. Starting on day 7, mice received 5×10^5^ freshly sorted ILC2 cells or vehicle 2 times per week for 4 weeks. s.c: subcutaneous; i.v.: intravenous. (B) Tumor volume in mice that received ILC2s (ILC2s; n=6) vs. mice that received vehicle (PBS; n=8). Data show mean ±sem. ns: not significant (Student t test).

**Supplementary Figure 4.** Immune cell composition in spleens of tumor-bearing mice. (A) Schematic representation of experimental design. 6-8-week old C56BL/6J mice were inoculated with 1×10^5^ LLc1 lung tumor cells on day 0, then partially lymphodepleted with CTX on day 6. Starting one day after CTX administration, mice received 5×10^5^ freshly sorted ILC2 cells or vehicle 2 times per week for 4 weeks. Mice were sacrificed 5 days after the last injection and spleens were collected. s.c: subcutaneous; i.p: intraperitoneal; i.v.: intravenous. (B) Frequency of tumor-infiltrating CD4 T cells identified as CD3^+^ CD4^+^, CD8 T cells identified as CD3^+^ CD8^+^ and CD4 regulatory T (Treg) cells identified as CD3^+^ CD4^+^ Foxp3^+^ in vehicle (CTX, n=5)- and ILC2-treated (CTX+ILC2, n=6) tumor-bearing mice. (C) Frequency of tumor-infiltrating conventional DCs (cDCs) identified as CD11c^+^ MHCII^+^ cells, granulocytic myeloid-derived suppressor cells (G-MDSCs) identified as CD3^-^ CD11b^+^ Ly6G^+^ Ly6C^low^, and monocytic MDSCs (M-MDSCs) identified as CD3^-^ CD11b^+^ Ly6G^-^ Ly6C^hi^ in vehicle (CTX, n=5)- and ILC2-treated (CTX+ILC2, n=6) tumor-bearing mice. Horizontal bars mark mean. Error bars mark sem. Statistical significance was calculated by unpaired two-tailed t-test. ns, not significant.

## Conflict of Interest

The authors declare that the research was conducted in the absence of any commercial or financial relationships that could be construed as a potential conflict of interest.

## Notes

### Competing Interest Statement

The authors have declared no competing interest.

