## Supplementary figures and images for "ILC2 cells promote lung cancer and accumulate in tumors concomitantly with immune-suppressive cells in humans and mice"

### Supplementary Figure 1

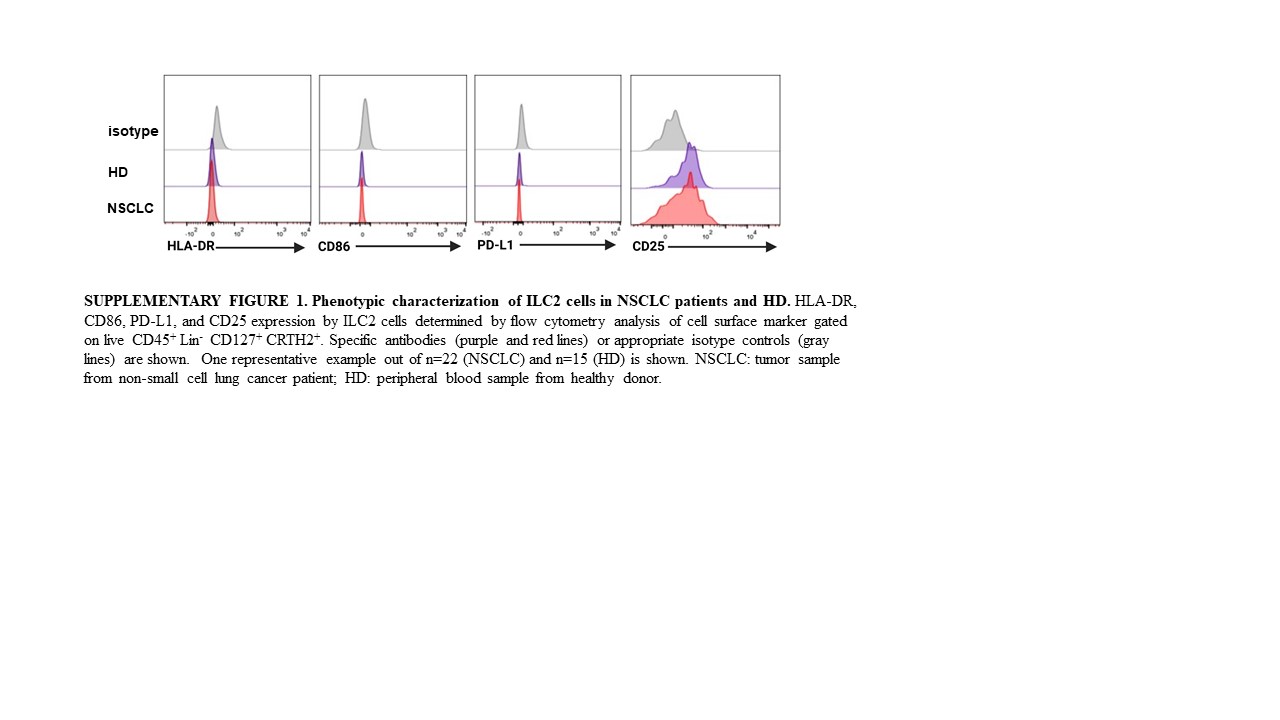

### Supplementary Figure 2

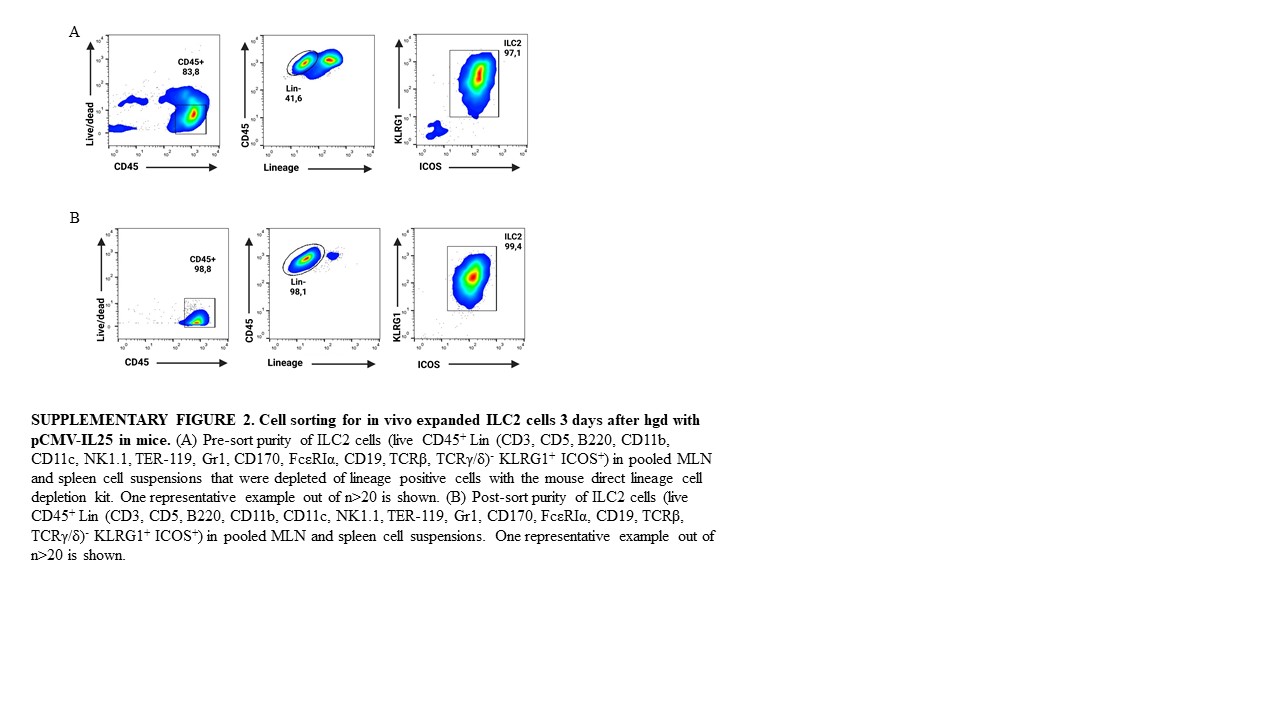

### Supplementary Figure 3

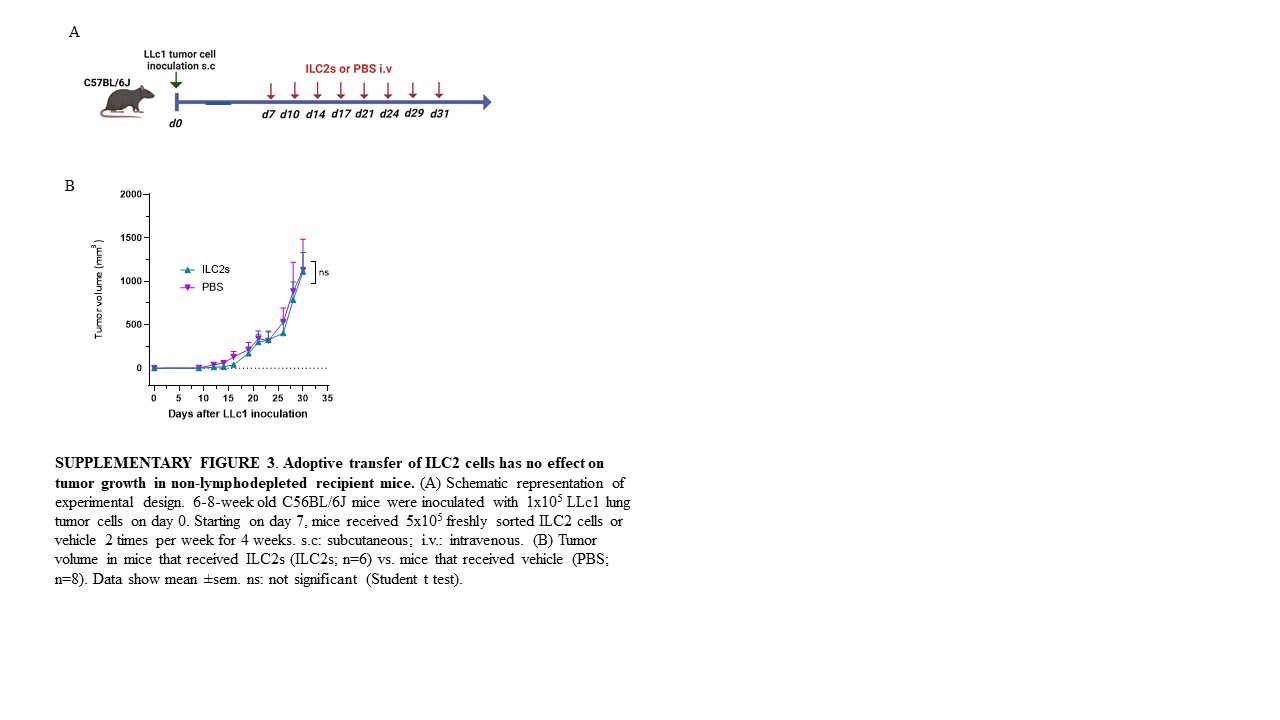

### Supplementary Figure 4

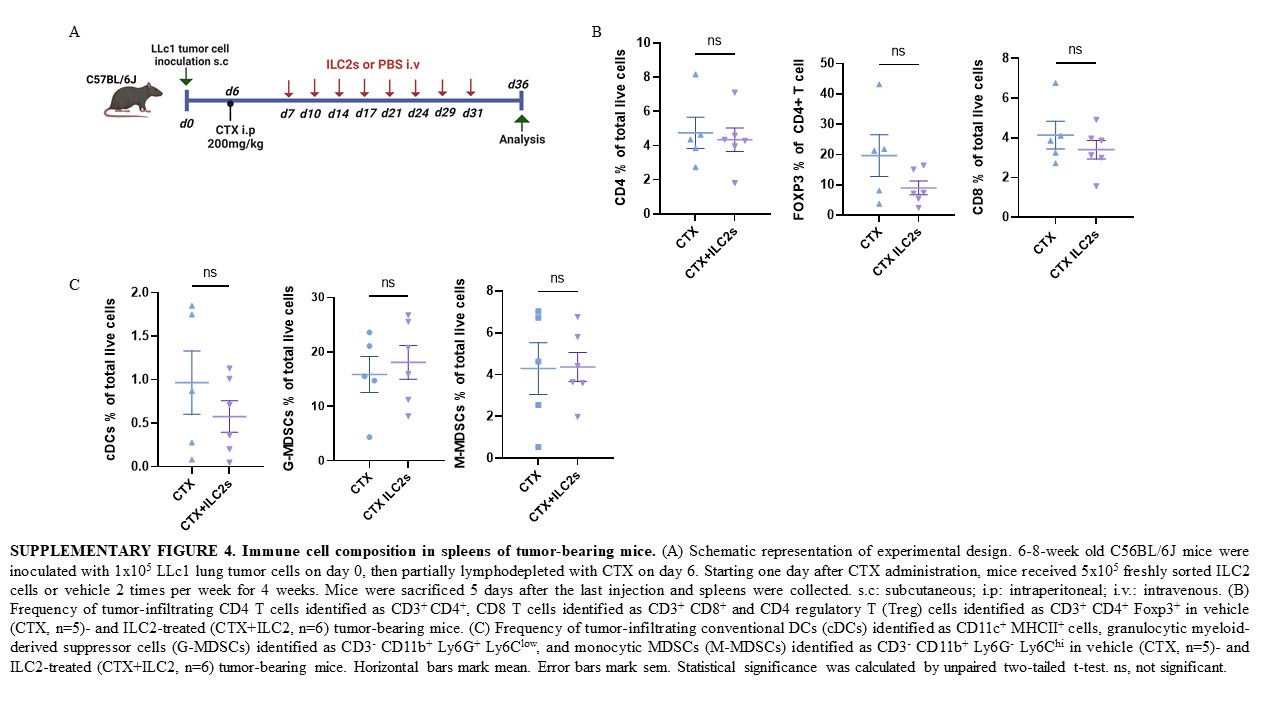
