## Supplementary Table 1 for "ILC2 cells promote lung cancer and accumulate in tumors concomitantly with immune-suppressive cells in humans and mice"

| **Patient characteristics** | |  | **NSCLC patients (n=28) n (%)** |
| --- | --- | --- | --- |
| Age | <65 |  | 20 (57,1 %) |
|  | >= 65 |  | 8 (22,9 %) |
| Gender | Female |  | 6 (17,1 %) |
|  | Male |  | 22 (62,9 %) |
| T Stage | Tx |  | ­ |
|  | T1 |  | 7 (25,0 %) |
|  | T2 |  | 10 (35,7 %) |
|  | T3 |  | 7 (25,0 %) |
|  | T4 |  | 4 (14,3 %) |
| N Stage | N0 |  | 14 (50,0 %) |
|  | N1 |  | 9 (32,1 %) |
|  | N2 |  | 5 (17,9 %) |
| M Stage | M0 |  | 27 (100 %) |
|  | M1 |  | ­ |
| Survival | Ex |  | 11 (61,1 %) |
|  | Alive |  | 7 (38,9 %) |
| Overall Survival (months) | Ex |  | 23,82 |
|  | Alive |  | 37 |
| Smoking Status | Active |  | 15 (53,6 %) |
|  | Quitted |  | 9 (32,1 %) |
|  | Non-Smoker |  | 4 (14,3 %) |

Supplementary Table S1. Clinicopathological characteristics of patients with NSCLC.
